# Cannabidiol prevents mucosal HIV-1 transmission by targeting Langerhans cells, dendritic cells, macrophages and T-cells

**DOI:** 10.64898/2026.01.07.697949

**Authors:** Caio César Barbosa Bomfim, Hugo Génin, Jammy Mariotton, Isabelle Matias, Daniela Cota, Nicolas Barry Delongchamps, Marc Zerbib, Morgane Bomsel, Yonatan Ganor

## Abstract

HIV-1 transmission depends on the structure and immune cell composition of mucosal epithelia. Transmission mechanisms involve direct infection of CD4^+^ T-cells or macrophages, and indirect viral transfer to CD4^+^ T-cells from Langerhans cells (LCs) or dendritic cells (DCs). LCs-mediated HIV-1 transfer is inhibited by the neuropeptide calcitonin gene-related peptide (CGRP), due to upstream activation in LCs of the transient receptor potential vanilloid 1 (TRPV1) ion channel. Herein, we investigated the potential anti-HIV-1 roles of cannabidiol (CBD), the non-psychoactive compound in *Cannabis sativa*, which has well-described immunosuppressive functions and principally activates TRPV1 over its cognate CB1 and CB2 receptors. We found that via TRPV1 activation, CBD inhibits *in-vitro* infection of mucosal HIV-1 cellular targets. Specifically, CBD inhibits macrophages HIV-1 direct infection, and CD4^+^ T-cells HIV-1 direct infection or upon viral transfer from LCs and DCs. Moreover, inhibition of macrophages infection and LCs-mediated HIV-1 transfer involves secreted CGRP. Importantly, CBD also blocks early events of HIV-1 transmission *ex-vivo* in human inner foreskin tissues, namely formation of epidermal LC-T-cell conjugates and resulting CD4^+^ T-cells infection. Altogether, CBD inhibits infection of all HIV-1 cellular targets, and commercial CBD products might be repositioned as novel HIV-1 pre-exposure prophylaxis, namely ‘CBD PrEP’.

## Introduction

The main route of HIV-1 entry and transmission is via mucosal epithelia of the male and female genital tracts during unprotected sexual intercourses. In the inner foreskin and vagina, incoming HIV-1 virions are rapidly internalized by resident antigen-presenting cells (APCs), including LCs and the recently identified epidermal DCs (EpiDCs), as we and others have shown ^1–6^. In turn, LCs and EpiDCs locally transfer infectious HIV-1 to CD4^+^ T-cells, leading to dissemination of the infection. Such transfer occurs in two phases, initiated by early transfer of virion escaping degradation, followed by late transfer of *de-novo* produced virions upon limited but productive LCs/EpiDCs HIV-1 infection ^7, 8^. Mucosal macrophages, as we reported in the penile urethra and others in the vagina, also support productive HIV-1 infection upon mucosal inoculation ^9–12^.

Protection against invasion by mucosal pathogens involves neuroimmune interactions between sensory peripheral nociceptor pain neurons innervating all mucosal epithelia, and resident immune cells ^13^. One central neuroimmune modulator is the nociceptor-derived mucosal neuropeptide CGRP ^14^, which we discovered to exert unexpected anti-HIV-1 activities. CGRP activates its cognate receptor expressed in LCs, and via several cooperative cellular and molecular mechanisms, strongly inhibits LCs-mediated HIV-1 transfer to CD4^+^ T-cells, but not direct HIV-1 infection of CD4^+^ T-cells ^15–21^. For instance, CGRP enhances LCs surface expression of atypical double-trimmer oligomers of the HIV-1-binding pathogen recognition lectin langerin ^20^, thereby limiting HIV-1 capture and diverting the virus from endolysosomal towards faster proteasomal degradation ^17^. CGRP also reduces LCs surface expression of adhesion molecules - thereby decreasing formation of LC-T-cell conjugates ^15, 19, 21^ - as well as the HIV-1 co-receptor CCR5, thus limiting their HIV-1 infection ^15, 16^. Together, these and other mechanisms, observed *in-vitro*, *ex-vivo* and *in-vivo*, inhibit LCs-mediated mucosal HIV-1 transmission.

Nociceptors are the main source of CGRP ^22^, which they secret following activation of their Ca^2+^ ion channel TRPV1. Activation of TRPV1 is also the mechanism enabling nociceptors to respond to noxious stimuli and generate action potentials, which are subsequently propagated to the central nervous system (CNS) and perceived as pain ^23^. We found that, like nociceptors, LCs express functional TRPV1 whose activation by its principal natural agonist capsaicin (CP), the spicy component in chili peppers, induces secretion of CGRP that inhibits LCs-mediated HIV-1 transmission ^21^. Moreover, CP-mediated TRPV1 activation in CD4^+^ T-cells inhibits their direct HIV-1 infection, but independently of CGRP ^21^. These results suggest potential benefit of CP against HIV-1, yet the suitability for such purpose of commercial CP-containing formulations, which induce an initial uncomfortable ‘flare’ response followed by TRPV1 desensitization and pain relief, remains to be determined. Instead, other natural TRPV1 agonists, such as cannabinoids (CBs), might be more appropriate.

CBs include a variety of molecules that are subdivided into three groups ^24^: 1) phyto CBs derived from the plant *Cannabis sativa* (marijuana), such as the non-psychoactive CBD and the psychoactive Δ^9^tetrahydrocannabinol (THC); 2) endogenous CBs (eCBs) that are synthesized from membrane lipid precursors and function as neurotransmitters, such as *N*-arachidonoylethanolamine (anandamide, AEA), its structurally related eCBs *N*-oleylethanolamine (OEA) and *N*-palmitoylethanolamine (PEA), and 2-arachidonoylglycerol (2-AG); 3) synthetic CB agonists and CB receptor antagonists.

CBs exert their functions by interacting with their cognate CB1 and CB2 receptors (termed herein CB1 and CB2 for simplicity), as well as other receptors. Importantly, some CBs have only low affinities for CB1 and/or CB2, but act as higher affinity agonists of TRPV1 ^25–27^. Accordingly: i) While CBD displays low affinities for CB1 and CB2 (Ki=4350->10,000nM and 2400->10,000nM, respectively) and acts as an antagonist / negative allosteric modulator / inverse agonist of both receptors, it activates TRPV1 at higher affinity (estimated Ki=1,000nM) compared to CB1/CB2; ii) AEA preferentially activates CB1 (Ki=61-543nM) and TRPV1 (estimated Ki=270nM) over CB2 (Ki=279-1940nM); iii) OEA and PEA do not activate CB1 and CB2; iv) Arachidonyl-2’-chloroethylamide (ACEA) is a synthetic agonist with high affinity for CB1 (Ki=1.4-5.3nM for CB1 vs. 195->2000nM for CB2) and dimethylbutyl-deoxy-Δ^8^-THC (JWH133) is a synthetic agonist with high affinity for CB2 (Ki=677nM for CB1 vs. 3.4nM for CB2).

The distribution of CB1 and CB2 in the human body is heterogenous. CB1 is primary expressed in the CNS and related to homeostatic function ^28^, and CB2 is mainly expressed in peripheral tissues, particularly in immune cells where it exerts immunosuppressive and anti-inflammatory functions ^29^. Due to such immunomodulatory functions, CBs have been mainly investigated for their potential to control chronic immune activation in the context of HIV-1 infection. In people living with HIV-1 (PLWH) under suppressive anti-retroviral therapy (ART), cannabis use is associated with decreased levels of T-cell activation, inflammatory monocytes and pro-inflammatory cytokines secretion ^30^, all of which are related to HIV-1 disease progression and comorbidities. Administration of oral CBs leads to significant reduction in surrogate markers of gut mucosal damage, systemic inflammation, and cellular immune activation, exhaustion, and senescence ^31^. Other studies *in-vitro* have indicated that THC and/or synthetic CB2 agonists reduce HIV-1 infection ^30^, e.g., in T-cells, macrophages and DCs ^32–35^.

Whether CBs, and especially CBD, could be useful for prevention of mucosal HIV-1 transmission, in addition to management of chronic HIV-1 infection, remains unclear. Herein, we investigated the capacity of CBD, as well as AEA, to inhibit the first steps of mucosal HIV-1 transmission and infection, using our established *in-vitro* and *ex-vivo* models of LCs/DCs-mediated HIV-1 transfer to CD4^+^ T-cells, and direct infection of CD4^+^ T-cells and macrophages. We also defined the distinct roles of TRPV1, secreted CGRP, and CB1/CB2 using specific agonists and antagonists.

## Materials and Methods

### Cells and tissues

PBMCs were separated from whole blood of healthy HIV-1-seronegative individuals by Ficoll gradient. CD14^+^ monocytes and CD4^+^ T-cells were isolated from PBMCs by negative magnetic selection (Stemcell Technologies). Monocytes (1x10^6^/well, in 12-wells plate) were differentiated into MDDCs or MDLCs ^36^ in complete medium (RPMI 1640 medium (Gibco Invitrogen) with 10% fetal bovine serum (Dominique Dutscher), 2mM L-glutamine (Gibco), 100U/ml penicillin (Gibco) and 100µg/ml streptomycin (Gibco)), supplemented with 100ng/ml granulocyte-macrophage colony-stimulating factor (GM-CSF) and 10ng/ml interleukin 4 (IL-4) for MDDCs, and additional 10ng/ml transforming growth factor beta 1 (TGF-β1) for MDLCs (all cytokines from Miltenyi Biotec). On days 2 and 4 of culture, 0.5ml of complete medium with the same cytokines at the same concentrations was added to each well, and cells were used on day 7 of culture. Monocytes were also differentiated into MDMs, by seeding into low attachment bacterial petri dishes (3x10^6^/dish), in complete medium supplemented with 50ng/ml GM-CSF and 25ng/ml macrophage colony-stimulating factor (M-CSF) for 7 days. A CD4^high^CCR5^high^ GFP-reporter T-cell line was a kind gift from Dr O. Kutsch (The University of Alabama at Birmingham) and was used we as described previously ^17^. All cells were maintained at 37°C and 5% CO_2_.

Normal foreskin tissues were obtained from healthy adults undergoing circumcision (Urology Service, Cochin Hospital, Paris), under informed consent and ethical approval (Comités de Protection des Personnes CPP Paris-IdF XI, N.11016). Cell suspensions were prepared by enzymatic digestion with dispase, followed by trypsin (for epidermal suspensions) or collagenase/DNase (for dermal suspensions), as we previously described ^1, 21^.

### Virus and infected cells

Viral stocks of the clade B HIV-1 molecular clones (NIH AIDS reagent program) ADA (R5), JRCSF (R5) or LAI (X4) were prepared by transfection of 293T cells, and of the HIV-1 primary isolate 93BR029 (V29) by amplification on PHA+IL-2-activated PBMCs. Stocks were quantified using the p24 Innotest HIV-1 ELISA (Fujirebio), according to the manufacturer’s instructions. PBMCs highly infected with HIV-1 V29 were prepared as we reported ^1^.

### Flow cytometry

MDLCs, MDMs, primary CD4^+^ T-cells (1x10^5^ cells/well) or inner foreskin cells (5x10^5^ cells/well) were stained for 30min (25μl/well) in round-bottom 96-well plates. Abs were diluted in phosphate buffer saline (PBS) and incubated at 4°C for surface staining, or using the Cytofix/Cytoperm kit (BD Biosciences) and incubation at room temperature for intracellular staining. The following fluorescently-conjugated Abs to human markers were used: pacific orange (PO) mouse-anti-CD45 (clone HI30, 1:200; Life Technologies); allophycocyanin (APC) mouse-anti-CD1a (clone HI149, 1:10; BD Pharmingen), mouse-anti-CD3 (clone Sk7, 1:10; BioLegend), mouse-anti-CD4 (clone SK3 1:5; BD) and mouse-anti-CD25 (clone 2A3, 1:5; BD); phycoerythrin (PE) recombinant human-anti-CD207 (clone REA770, 1:100; Miltenyi), mouse-anti-CD8 (clone HIT8a, 1:10; BD) and mouse-anti-HIV-1 p24 (clone KC57, 1:160; Beckman Coulter); AlexaFluor 488 (AF488) mouse-anti-CD4 (clone SK3 1:10; BioLegend) and mouse-anti-CB1 (clone 368302, 1:20; R&D Systems); fluorescein isothiocyanate (FITC) mouse-anti-CCR5 (clone 2D7, 1:10; BD), mouse-anti-CD69 (clone FN50, 1:50; BD) and rabbit-anti-CB2 (Item No. 10010712, 1:40; Cayman Chemical). Negative control included staining with matched isotype Abs or fluorescence minus one (F-1). When indicated, viability was determined using the Viobility Fixable Dye staining kit (Miltenyi), according to the manufacturer’s instructions. Samples were acquired using a Guava easyCyte flow cytometer (Merck-Millipore) and analyzed with FlowJo software (FlowJo LLC).

### Western blot

MDLCs (2x10^6^) were lysed for 30min on ice with 100µl lysis buffer (50mM Tris buffer pH=7.5, 150nM NaCl, 2mM EDTA, 1% Triton X-100, 0.1% SDS, 1:100 dilutions of phosphatase inhibitors II/III and protease inhibitor cocktail (Sigma)), followed by three cycles of 10sec vortex and 10min incubation on ice. Lysates were centrifuged for 10min at 4°C/13,200 rpm, and supernatants were collected and stored at −80°C. Protein contents in cell lysates were quantified using the BCA kit (ThermoFisher) according to the manufacturer’s instructions, and 20µg proteins were mixed with loading buffer (100mM Tris buffer pH=7.5, 5% β-mercaptoethanol, 12% glycerol, 5mM EDTA, 5% SDS, 0.01% bromophenol blue), heated for 5min at 95°C, run over a 10% SDS-PAGE, and transferred at 4°C onto nitrocellulose membranes. Blocking was performed for 1h at room temperature with blocking buffer (Tris-buffered saline (TBS), 0.5% Tween-20 and 0.5% dry milk). The blots were next incubated overnight at 4°C with 1μg/ml of rabbit IgG against human CB2 (#Cat PA1-744; Invitrogen), followed by 1:1000 dilution of horseradish peroxidase (HRP)-conjugated donkey-anti-rabbit IgG Ab (Southern Biotech) for 1h at room temperature. Loading control was verified by incubation with mouse IgG against GAPDH (1:1000; Santa Cruz,), followed by 1:5000 dilution of HRP-conjugated donkey anti-mouse IgG (Caltag). All Abs were diluted in blocking buffer, and pre-stained SDS-PAGE standard markers (ThermoFisher) were applied to determine molecular weights. Revelation was performed for 1-10sec with ECL-Prime chemiluminescence detection kit (Amsersham). Images were acquired with the Fusion FX camera platform (Vilver Lournmat) and protein expression was quantified with ImageJ software (NIH).

### LCs content of eCB

MDLCs (5-10x10^6^/sample) were left non-infected or infected with HIV-1 R5 ADA (250ng p24 per 10^6^ cells) for 24h at 37°C. Cells were next washed, fixed with methanol for 20min at room temperature, centrifuged, and the cell pellets stored at −80°C. The levels of different eCBs were next determined following cell pellet homogenization, extraction and analysis by LC-MS/MS, as we described ^37^. The amounts of eCBs were expressed as pmols per total mg of proteins extracted from the fixed cells.

### HIV-1 transfer and infection

MDLCs or MDDCs (0.5-1x10^5^/well) were pre-treated for 24h at 37°C with the indicated concentrations of CBD, AEA, ACEA, JWH133 (all from Tocris Biotechne) or CGRP (Sigma), in triplicates in a round-bottom 96-wells plate (200µl/well final). When indicated, the TRPV1 antagonist A425619 (10^-6^M; Sigma) or the CGRP receptor antagonist BIBN (10^-6^M; Sigma) were added 15min before agonists. Of note, we verified that at the concentrations used herein, none of the agonists and antagonists affected cell viability (>90%, data not shown). Next, cells were washed and pulsed with HIV-1 R5 ADA (5ng or 25ng p24 per well, for cumulative or temporal transfer assays, respectively) for 4h at 37°C. The cells were then washed, and co-cultured with autologous CD4^+^ T-cells (3x10^5^/well, added immediately after viral pulse) or CD4^high^CCR5^high^ GFP-reporter T-cells (0.5x10^5^/well, added immediately or 48h after viral pulse). HIV-1 transfer was evaluated a week later by measuring p24 content in the co-culture supernatants using p24 ELISA (cumulative transfer), or by evaluating 48h later GFP fluorescence by flow cytometry (temporal transfer).

Primary blood CD4^+^ T-cells were activated for 48h with 5µg/ml PHA (Sigma) + 10U/ml IL-2 (Roche), followed by treatment for 24h at 37°C with the indicated concentrations of agonists. Cells were then pulsed with HIV-1 R5 JRCSF or X4 LAI (25ng or 10ng p24 per well, respectively) for 4h at 37°C and washed. After 4 days, cells were fixed and permeabilized with the Cytofix/Cytoperm kit (BD), stained for 30min at room temperature with PE-conjugated mouse-anti-HIV-1 p24 (see above), and examined by flow cytometry.

MDMs were treated for 24h at 37°C with agonists (10^-6^M). When indicated, the CGRP receptor antagonist BIBN (10^-6^M) was added 15min before agonists. The cells were then pulsed for 4h with HIV-1 R5 ADA (25ng p24 per well), washed, further cultured for 7 days, and HIV-1 p24 content was measured in the supernatants by p24 ELISA.

### CGRP secretion

MDLCs (1x10^6^/well) were treated for 24h at 37°C with the indicated concentrations of agonists, in a round-bottom 96-wells plate (200µl/well final). The culture supernatants were collected immediately following treatment and CGRP levels were measured using a competitive CGRP enzyme immunoassay kit (EIA; Phoenix Pharmaceuticals) according to the manufacturer’s instructions.

### *Ex-vivo* infection of inner foreskin tissue explants

Round inner foreskin tissue pieces were cut using a Harryis Uni-Core of 8-mm diameter, transferred to a 24-wells plate (in duplicates) and incubated submerged for 24h at 37°C in 1ml complete RPMI medium, alone or supplemented with CBD or AEA (10^-5^M). The media were next collected and the levels of secreted CGRP were measured with the CGRP EIA mentioned above. Next, the explants were washed, transferred to two-chamber Transwell inserts with polycarbonate membranes (Costar), and cloning ring cylinders were adhered to the apical surface using surgical glued. HIV-1 was next inoculated in a polarized manner for 4h at 37°C, by adding either non-infected or HIV-1 V29-infected PBMCs to the inner space of cylinders, as we previously described and recently optimized ^1, 21^. Epidermal cell suspensions were prepared immediately after inoculation, and dermal cell suspensions were prepared following washing of the explants, additional incubation for three days at 37°C submerged in 1ml fresh medium, and subsequent enzymatic digestion, as we described above. Pooled cells of each duplicate were resuspended in PBS, transferred to a round-bottom 96-wells plate, stained and examined by flow cytometry as described above.

### Statistical Analyses

Data was analyzed using Prism software (GraphPad). Concentration-response curves were analyzed with the [log(inhibitor) vs. normalized response - variable slope] model for HIV-1 trans-infection inhibition, with the -log molar concentrations of agonists generating 50% response representing potencies (i.e. pIC50). For comparing secreted CGRP levels from matched tissue explants, pairwise comparisons were performed using the unpaired non-parametric Mann-Whitney test. The remaining statistical significance was analyzed with the two-tailed Student’s t-test. Differences between groups were considered significant when the p-values were <0.05.

## Results

### LCs express CB1 and CB2 and produce eCBs

CB2 is mainly expressed in immune cells, with some studies reporting a hierarchical pattern of expression (i.e., B-cells > natural killer cells > monocytes > T-cells ^38^). Yet, the precise levels of CB2 expression in different immune cell populations remain controversial. For instance, some studies showed that human monocytes do not express surface CB2 ^39, 40^ while others could detect the presence of human CB2^+^ monocytes ^41^. In addition, while CB1 expression is mainly detected in the CNS, functional CB1 is also expressed by immune cells ^42^.

Based on our previous study demonstrating CB1 and CB2 expression in human DCs ^43^, we evaluated their expression also in human LCs. We isolated CD14^+^ monocytes by negative magnetic selection from peripheral blood mononuclear cells (PBMCs), and differentiated the cells into monocyte-derived LCs (MDLCs) as previously described ^36^. Using antibodies (Abs) directed against extracellular epitopes of CB1 and CB2 followed by flow cytometry, we detected surface expression of CB1 and CB2 on langerin^+^ MDLCs, with CB2 expressed at much lower levels than CB1 (Figure 1A). To further confirm that MDLCs indeed express CB2, we evaluated its intracellular levels and found that the majority of MDLCs expressed high CB2 levels in intracellular compartments (Supplementary Figure 1A), in line with the reported elevated levels of intracellular CB2 protein stores in different PBMCs subpopulations, including monocytes and T-cells ^39^. Such intracellular CB2 has been shown to be functional in different cell types, including immune cells ^44^. Western blot (WB) analysis also demonstrated clear CB2 expression in MDLCs (Figure 1B).

**Figure 1.**
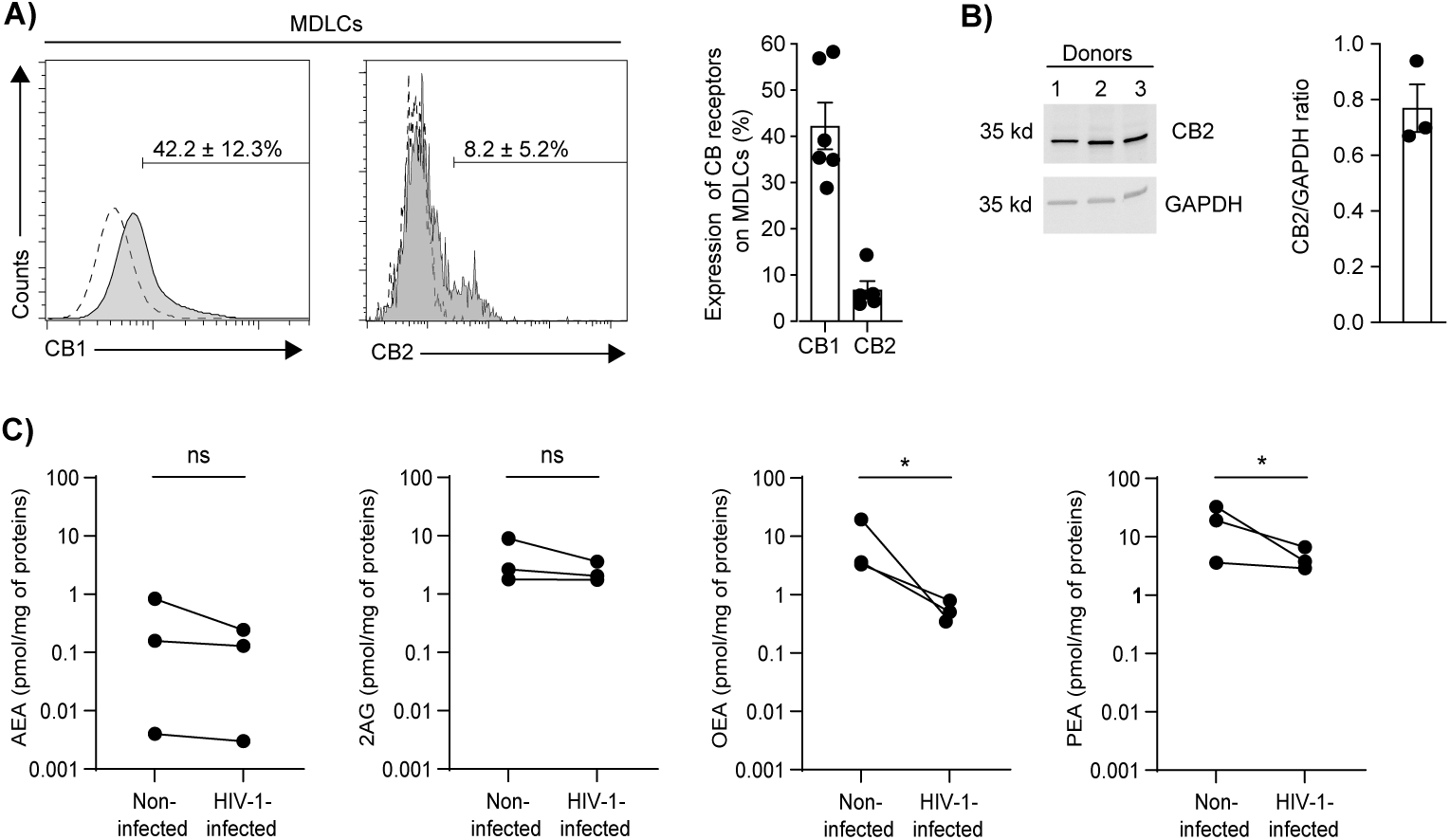
CB1 and CB2 expression and eCBs production in human LCs. **(A)** Representative flow cytometry overlay histograms showing CB1 and CB2 surface expression in MDLCs, gated on langerin^+^ cells. Numbers and graph show mean±SD percentages of CB1^+^ or CB2^+^ cells. **(B)** CB2 protein levels detected by WB in total cell lysates of MDLCs and quantified by densitometry. Shown is a representative blot, and graph shows mean±SD CB2/GAPDH ratios. **(C)** AEA, 2-AG, OEA and PEA levels evaluated by LC-MS/MS, in cell lysates of non-infected MDLCs vs. MDLCs infected with HIV-1 R5 ADA for 24h, and expressed as pmol per mg of total proteins. In all graphs, dots denote different experiments using cells prepared from different human donors; *p<0.05, Student’s t-test.

Different from other neurotransmitters that are pre-stored intracellularly before release, eCBs are lipids and cannot be stored in vesicles; their biosynthesis is induced from plasma membrane lipid precursors when needed (‘on-demand’). A variety of immune cells synthetize eCBs, including human DCs as we reported ^43^. To determine whether human LCs also produce eCBs, we subjected cell lysates of MDLCs to high-performance liquid chromatography - mass spectrometry (LC-MS/MS). We quantified the levels of AEA and 2-AG, the two most studied eCBs, as well as that of the two eCB-like OEA and PEA (i.e., structurally related to AEA) that have no agonistic activity against CB1 and CB2 but potentiate AEA actions via TRPV1 ^45^. Such measurements were performed comparatively, using cell lysates prepared from non-infected MDLCs vs. MDLCs that were infected for 24h with the HIV-1 laboratory strain ADA of R5 tropism (i.e., as sexually transmitted HIV-1 is predominantly R5-tropic). These experiments showed that all four eCBs were detected in untreated MDLCs, with AEA found at the lowest levels (Figure 1C), as we found in human DCs ^43^ and others in T-cells ^46^. Interestingly, HIV-1 infection of MDLCs decreased the levels of these eCBs, with statistically significant reduction for OEA and PEA (Figure 1C).

These results show that LCs express CB receptors and produce eCBs. The data also suggest that HIV-1 has evolved mechanisms to counteract eCBs, indicative of their potential anti-viral roles.

### CBD and AEA inhibit HIV-1 R5 transfer from LCs and DCs to CD4^+^ T-cells

Upon its mucosal entry, HIV-1 is rapidly internalized by LCs that degrade the majority of virions. However, some intact and infectious virions are retained within LCs and can be transferred to CD4^+^ T-cells ^8^. To evaluate whether CBs play any role during HIV-1 transfer, we used our previously described HIV-1 transfer assay ^15^ (Figure 2A). Accordingly, MDLCs were left untreated or pre-treated for 24h with a range of concentrations of CBD, AEA, the CB1 agonist ACEA, the CB2 agonist JWH133, and also CGRP as positive control. Of note, JWH133 is a lipophilic molecule that crosses the plasma membrane and reaches intracellularly located CB2 ^47^. MDLCs were then washed, pulsed for 4h with HIV-1 R5 ADA, washed again, and co-cultured with autologous CD4^+^ T-cells. HIV-1 replication in CD4^+^ T-cells, which we have previously shown to result from MDLCs-mediated viral transfer ^15^, was measured in the co-culture supernatants a week later by p24 ELISA.

**Figure 2.**
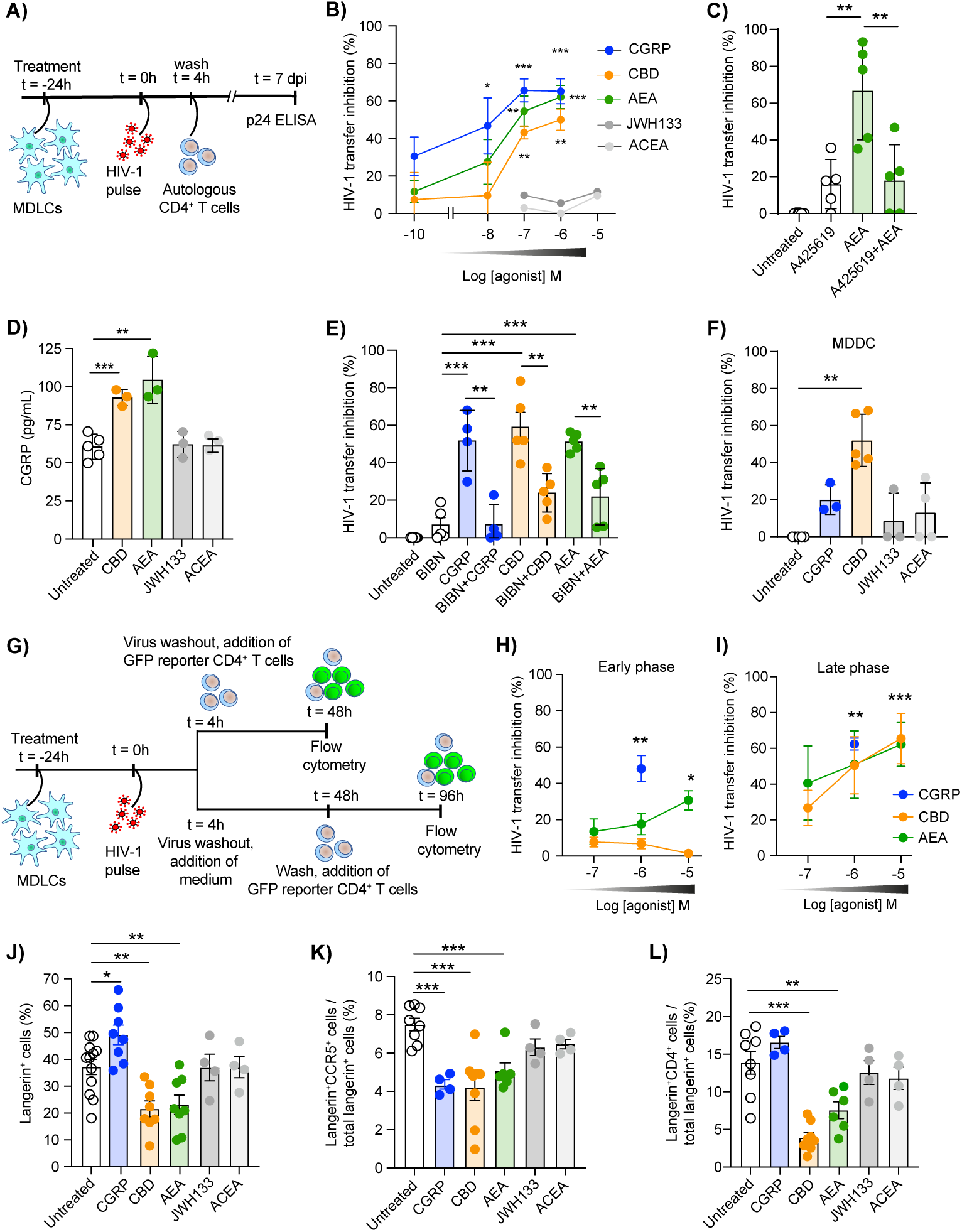
CBD and AEA inhibit HIV-1 R5 transfer from human LCs and DCs to CD4^+^ T-cells. **(A)** Graphical representation of the assay evaluating HIV-1 transfer from MDLCs to autologous CD4^+^ T-cells, cumulating the early and late phases of transfer. **(B)** MDLCs were left untreated or pre-treated for 24h with the indicated molar concentrations of CBD, AEA, ACEA, JWH133 or CGRP, pulsed with HIV-1 R5 ADA for 4h, and washed. Cells were then co-cultured with autologous CD4^+^ T-cells, and the supernatants were evaluated 7 days post infection (dpi) for p24 content by ELISA. Graph shows mean±SD (n=3-9) percentages of HIV-1 transfer inhibition, calculated against untreated cells serving as set point. **(C)** MDLCs were left untreated or pre-treated for 24h with AEA (10^-6^M), in the absence or presence of the TRPV1 antagonist A425619 (10^-6^M) added 15min before AEA. HIV-1 transfer was then evaluated as above, and graph shows mean±SD percentages of HIV-1 transfer inhibition. **(D)** MDLCs were left untreated or pre-treated for 24h with agonists (10^-5^M). The cells were then washed, cultured in fresh medium for additional 48h, and secretion of CGRP into the culture supernatants was evaluated using a competitive CGRP EIA. Graph shows mean±SD levels of pg/ml secreted CGRP. **(E)** MDLCs were pre-treated for 24h with CBD, AEA or CGRP (10^-6^M), in the absence or presence of the CGRP receptor antagonist BIBN (10^-6^M) added 15min before agonists. HIV-1 transfer was then evaluated and is presented as above. **(F)** MDDCs were pre-treated for 24h with CBD, ACEA, JWH133 (10^-5^M) or CGRP (10^-6^M), and HIV-1 transfer was then evaluated and is presented as above. **(G)** Graphical representation of the assay evaluating temporally the early and late phases of HIV-1 transfer from MDLCs to GFP-reporter T-cells. **(H, I)** MDLCs were left untreated or pre-treated for 24h with the indicated molar concentrations of CBD, AEA or CGRP, pulsed with HIV-1 R5 ADA for 4h, and washed. Cells were then co-cultured with CD4^high^CCR5^high^ GFP-reporter T-cells, added immediately after pulse (H) or 48h later (I), followed by flow cytometry 48h later. Graph show mean±SD (n=3-8) percentages of HIV-1 transfer inhibition, calculated against untreated cells serving as set point. **(J-L)** MDLCs were left untreated or pre-treated for 24h with agonists (10^-5^M) or CGRP (10^-6^M), followed by flow cytometry. Graphs show mean±SD percentages of positive cells expressing langerin (J), or proportions (out of langerin^+^ cells) of cells expressing CCR5 (K) or CD4 (L). In all graphs, dots denote different experiments using cells prepared from different human donors; *p<0.05, **p<0.005 and ***p<0.0005, Student’s t-test.

These experiments revealed that both CBD and AEA inhibited HIV-1 transfer in a dose-depended manner, as did CGRP that was the most potent (Figure 2B), with calculated potencies (expressed as -log of the half-maximal inhibitory concentrations (pIC_50_) [95% confidence intervals; R^2^]) of 6.18 [7.08-0.51; 0.87] for CBD; 6.91 [7.50-6.09; 0.96] for AEA; and 7.96 [9.08-7.04; 0.93] for CGRP. In contrast, ACEA and JWH133 had no effects on MDLCs-mediated HIV-1 transfer (Figure 2B), suggesting that CBD and AEA inhibit HIV-1 transfer by activating TRPV1 and inducing CGRP release.

To verify this hypothesis, we first added the TRPV1 antagonist A425619 as before ^21^, 15min before pre-treating MDLCs with AEA (i.e., AEA activates both TRPV1 and CB1), and evaluated HIV-1 transfer as above. These experiments showed that A425619 alone had no significant effect, but fully abrogated AEA-mediated inhibition of HIV-1 transfer that was reduced to the levels of A425619 alone (Figure 2C). Second, treatment of MDLCs with both CBD and AEA increased CGRP secretion, while ACEA and JWH133 had no effects (Figure 2D). Third, we added the CGRP receptor antagonist BIBN4096 (BIBN) as before ^21^, 15min before pre-treating MDLCs with agonists, and found that BIBN significantly and completely abrogated CGRP-mediated inhibition as expected, and approximately half of that mediated by CBD or AEA (Figure 2E).

In other experiments, we evaluated HIV-1 transfer using monocyte-derived DCs (MDDCs) instead of MDLCs. These experiments confirmed that CGRP had minimal effects as we previously reported ^15^, and further revealed that CBD significantly inhibited HIV-1 transfer mediated by MDDCs while JWH133 and ACEA had no significant effects (Figure 2F). LCs-mediated HIV-1 transfer is divided into two steps: the first early phase is mediated by non-degraded intact virions retained intracellularly and/or on the surface, and the second late phase is mediated by *de-novo* produced virions ^8^. As the assay described above models the whole HIV-1 transfer process, cumulating the first and second phases together, we determined which phase of HIV-1 transfer is affected by CBs using our temporal assay of HIV-1 transfer ^17^ (Figure 2G). Accordingly, MDLCs were left untreated or pre-treated for 24h with a range of concentrations of CBD, AEA, ACEA, JWH133 or CGRP. The cells were then pulsed with HIV-1 R5 ADA, washed, and co-cultured for 48h with CD4^high^CCR5^high^ green fluorescent protein (GFP)-reporter T-cells, followed by flow cytometry. The reporter cells were added to MDLCs either immediately or 48h following HIV-1 pulse, i.e. experimental settings that model the early or late phases of HIV-1 transfer, respectively (Figure 2G). These experiments showed that while CGRP inhibited both transfer phases as we previously reported ^17^, CBD and AEA inhibited the late phase, with no effect of CBD and only modest effect of AEA at its highest concentration tested on the early phase of HIV-1 transfer (Figure 2H and I).

To characterize the mechanisms by which CBD and AEA inhibit LCs-mediated late transfer of HIV-1 R5, we next tested for alterations in surface expression of langerin (i.e., permitting HIV-1 internalization and facilitating both early and late phases of HIV-1 transfer ^8^) and CD4/CCR5 (i.e., the HIV-1 receptor/co-receptor that facilitate productive infection and late HIV-1 R5 transfer). Of note, our previous studies showed that in MDLCs, CGRP increases langerin and decreases CCR5, while TRPV1 activation by CP does not affect langerin but decreases CCR5 ^15, 16, 19, 21^.

These experiments confirmed that CGRP increased langerin (Figure 2J) and decreased CCR5 (Figure 2K) as before, and further revealed that it had no effect on surface expression of CD4 (Figure 2L). In addition, CBD and AEA reduced CCR5 surface expression similarly to CGRP (Figure 2K), but also decreased surface expression of both langerin and CD4 (Figure 2J and L). In contrast, ACEA and JWH133 had no effects on expression of these receptors (Figure 2J-L). These results show that CBD and AEA inhibit LCs-mediated late HIV-1 transfer via TRPV1 activation leading to reduced langerin, CCR5 and CD4 expression. CCR5 decrease could be induced by CGRP released upon TRPV1 activation, while langerin and CD4 decrease is TRPV1-dependend but CGRP-independent. CBD also inhibits DCs-mediated HIV-1 transfer, in TRPV1-dependend but CGRP-independent manners.

### CBD, but not AEA, inhibits direct infection of CD4^+^ T-cells with HIV-1 R5

Previous studies reported that synthetic agonists of CB1 and/or CB2 decrease direct infection of CD4^+^ T-cells with HIV-1 of X4, but not R5, tropism ^33, 48^. In addition, AEA was reported to either have no effect ^49^ or inhibit CD4^+^ T-cells infection with HIV-1 X4 via CB2-mediated decreased CXCR4 activation ^48^. We recently demonstrated that the TRPV1 agonist CP, but not CGRP, blocks CD4^+^ T-cells infection with HIV-1 R5 ^21^. The effects of CBD remained to be determined.

Using flow cytometry as above, we first confirmed the surface expression of CB1 and CB2 in primary blood CD4^+^ T-cells (Figure 3A), which also expressed CB2 intracellularly (Supplementary Figure 1B).

**Figure 3.**
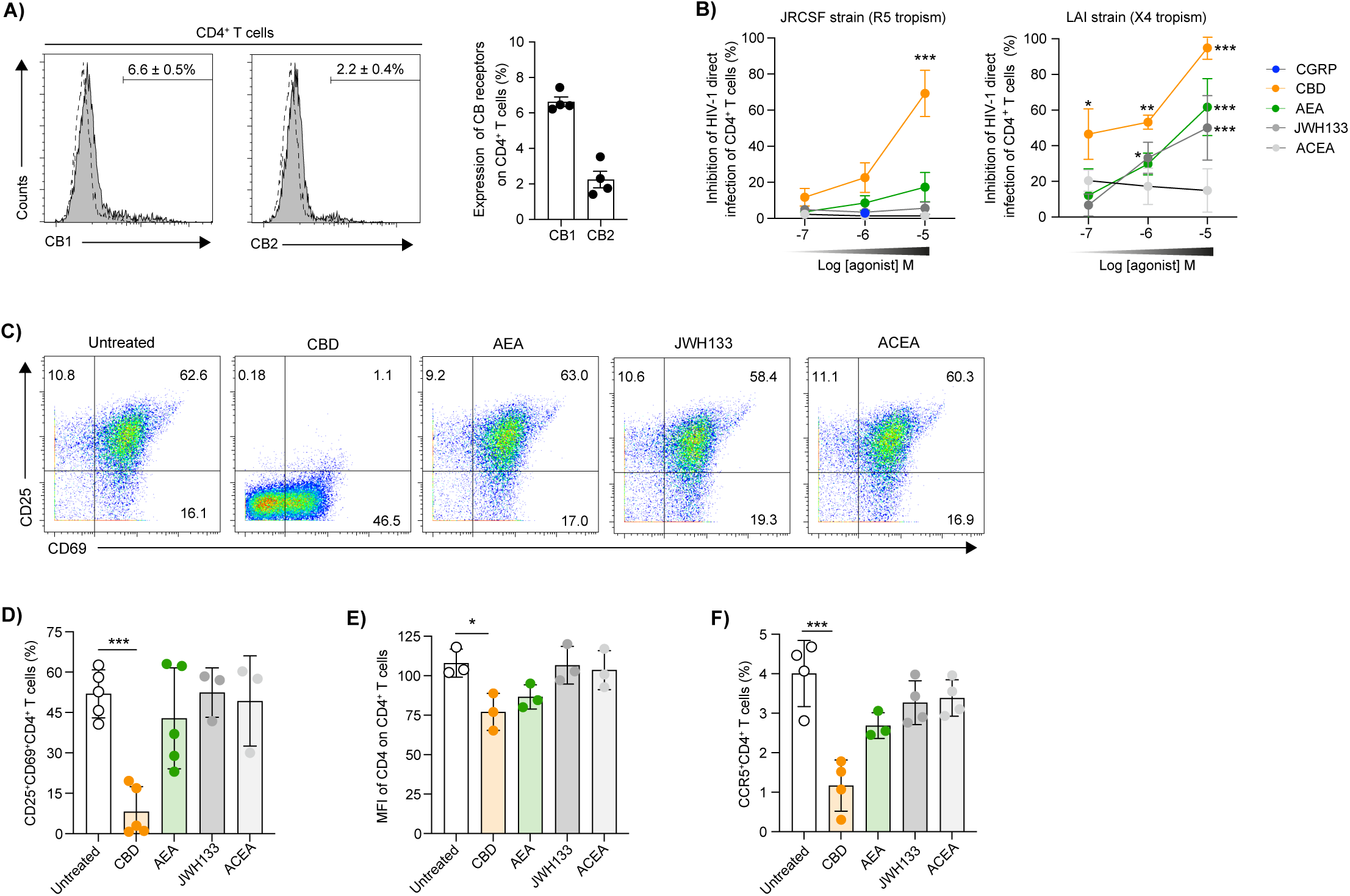
CBD, but not AEA, inhibits HIV-1 R5 direct infection of human CD4^+^ T-cells. **(A)** Representative flow cytometry overlay histograms showing CB1 and CB2 surface expression in primary blood CD4^+^ T-cells. Numbers and graph show mean±SD percentages of CB1^+^ or CB2^+^ T-cells. **(B)** PHA+IL-2-activated primary blood CD4^+^ T-cells were pre-treated for 24h with the indicated molar concentrations of CBD, AEA, JWH133, ACEA or CGRP. The cells were then washed, pulsed with HIV-1 R5 JRCSF or X4 LAI, and HIV-1 replication was evaluated 4 days later by p24 intracellular staining and flow cytometry. Graphs show mean±SD (n=4-6) percentages of HIV-1 infection inhibition, normalized against untreated cells serving as set point. **(C, D)** Primary blood CD4^+^ T-cells were activated for 48h with PHA+IL-2, alone or in the presence of CBD, AEA, JWH133 or ACEA (10^-5^M). The cells were then evaluated by flow cytometry for surface expression of the activation marker CD69 and CD25. In (C) shown are representative flow cytometry dot plots. In (D), graph shows mean±SD percentages of CD69^+^CD25^+^ double-positive CD4^+^ T-cells. **(E, F)** Primary blood CD4^+^ T-cells were left untreated or treated for 24h with agonists (10^-5^M), followed by flow cytometry evaluating surface expression of CD4 and CCR5. Graphs show mean±SD of CD4 MFI (E) or percentages of CCR5^+^CD4^+^ T-cells (F). In all graphs, dots denote different experiments using cells prepared from different human donors; *p<0.05, **p<0.005 and ***p<0.0005, Student’s t-test.

Next, to study the effect of CBs on direct HIV-1 infection of T-cells, primary blood CD4^+^ T-cells were activated with phytohemagglutinin (PHA) + interleukin 2 (IL-2) for 48h and then treated for additional 24h with a range of concentrations of CBD, AEA, ACEA, JWH133 or CGRP. The cells were next pulsed with HIV-1 laboratory strains, either R5-tropic JRCSF or X4-tropic LAI, and HIV-1 replication was evaluated 4 days later by p24 intracellular staining and flow cytometry.

As we reported, CGRP had no effect on HIV-1 R5 infection of CD4^+^ T-cells (Figure 3B). In addition, these experiments showed that CBD inhibited in a dose-dependent manner infection with both HIV-1 R5 and X4 (Figure 3B). In contrast, AEA and the synthetic agonists of CB1 and CB2 had no effects on infection with HIV-1 R5, yet AEA and the synthetic CB2 agonist JWH133 inhibited in a dose-dependent manner infection with HIV-1 X4 (Figure 3B) as reported ^33, 48^.

Activation of primary CD4^+^ T-cells is a pre-requisite for their infection with HIV-1 *in-vitro*. We therefore added CBD, AEA, ACEA or JWH133 during the PHA+IL-2 activation period and determined surface expression of the activation markers CD69 and CD25 by flow cytometry. These experiments showed that CBD, but none of the other agonists, significantly and strongly decreased the proportion of CD69^+^CD25^+^ double-positive CD4^+^ T-cells (Figure 3C and D). CBD-mediated inhibition of infection with HIV-1 R5 also correlated with decreased surface expression of CD4 (evident by lower mean fluorescent intensity (MFI); Figure 3E), and especially of CCR5 (evident by lower percentages of positive cells; Figure 3F). In contrast, AEA, ACEA and JWH133 had no effects on CD4 and CCR5 expression (Figure 3E and F). These findings show that by activating TRPV1, and independently of CGRP and CB1/CB2, CBD impairs T-cell activation and decreases their CD4/CCR5 expression, thereby limiting direct HIV-1 R5 infection of CD4^+^ T-cells.

### CBD, AEA and CGRP inhibit infection of macrophages with HIV-1 R5

A previous study reported that pre-treatment of human macrophages with CBD increases their infection with HIV-1 X4, while long-term presence of CBD following viral exposure results in decreased HIV-1 infection ^50^. Another study showed that CB2 activation blocks macrophage infection with HIV-1 R5 ^51^. We further studied the roles of CBD and AEA during infection of macrophages with HIV-1 R5-tropic.

Using human blood CD14^+^ monocytes differentiated into monocytes-derived macrophages (MDMs) as we previously described ^52^, we confirmed by flow cytometry that MDMs expressed surface CB1 and CB2 (Figure 4A), and also intracellular CB2 (Supplementary Figure 1C).

**Figure 4.**
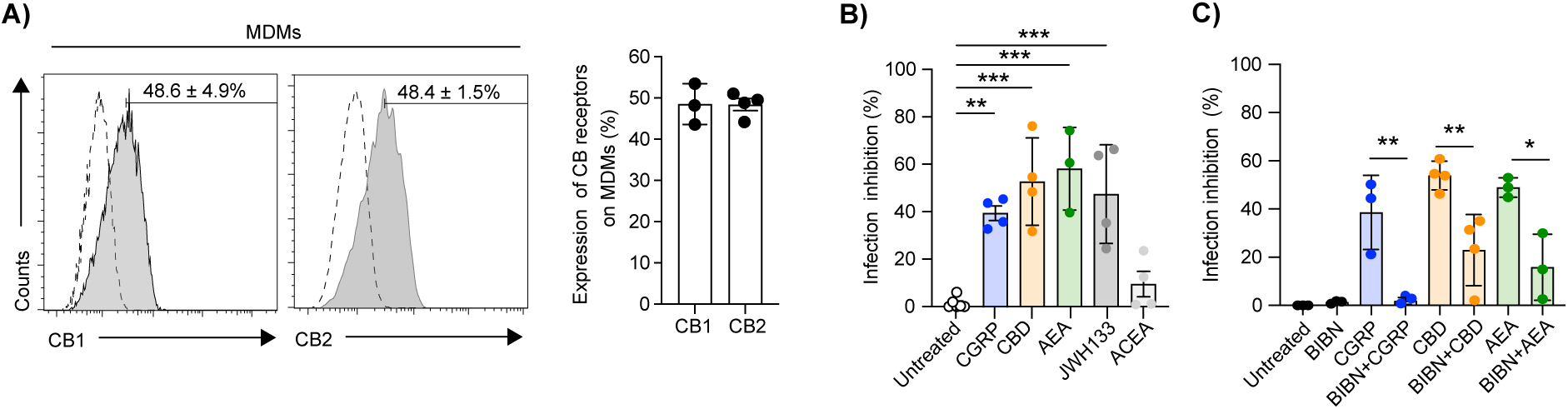
CBD, AEA, JWH133 and CGRP inhibit MDMs infection with HIV-1 R5. **(A)** Representative flow cytometry overlay histograms showing CB1 and CB2 surface expression in MDMs. Numbers and graph show mean±SD percentages of CB1^+^ or CB2^+^ MDMs. **(B, C)** MDMs were left untreated or pre-treated for 24h with CBD, AEA, JWH133, ACEA or CGRP (10^-6^M). The CGRP receptor antagonist BIBN (10^-6^M) was added 15 min before addition of agonists. The cells were then washed, pulsed with HIV-1 R5 ADA for 4h, washed again, and HIV-1 content in the culture supernatants was evaluated at 7 dpi by p24 ELISA. Graphs show mean±SD percentages of HIV-1 infection inhibition, normalized against untreated cells serving as set point. In all graphs, dots denote different experiments using cells prepared from different human donors; *p<0.05, **p<0.005 and ***p<0.0005, Student’s t-test.

Next, MDMs were pre-treatment for 24h with 10^-6^M of CBD, AEA, ACEA, JWH133 or CGRP, pulsed with HIV-1 R5 ADA, and HIV-1 content in the supernatants was examined 7 days later by p24 ELISA. These experiments revealed that both CBD and AEA decreased HIV-1 R5 infection of MDMs, as did the CB2 agonist JWH133 but not the CB1 agonist ACEA (Figure 4B). We also observed a significant reduction of MDMs infection following pre-treatment with CGRP, indicating that activation of the CGRP receptor could play a role in containment of HIV-1 infection in MDMs (Figure 4B). In agreement, addition of the CGRP receptor antagonist BIBN 15min before pre-treatment with agonists, significantly and completely abrogated the inhibitory effects of CGRP (Figure 4C) and approximately half of that induced by CBD and AEA (Figure 4C).

These results indicate that CBD and AEA inhibit HIV-1 R5 infection of macrophages, partially via activating TRPV1 and inducing CGRP secretion. Activation of CB2 also results in inhibition of HIV-1 R5 infection in MDMs.

### CBD inhibits mucosal HIV-1 R5 transmission *ex-vivo*

To extend the *in-vitro* findings described above, we prepared cell suspensions from human inner foreskin epidermal sheets as we established ^15^ and evaluated surface expression of CB1 and CB2 by flow cytometry. We distinguished between inner foreskin LCs and EpiDCs based on their reported differential expression of CD1a and langerin (CD207) ^6^, and confirmed as before the presence of CD1a^hi^CD207^hi^ LCs and CD1a^low^CD207^low/neg^ EpiDCs ^21^ in gated viable CD45^+^ cells (Figure 5A). We found that inner foreskin LCs expressed comparable levels of CB1 and CB2 (Figure 5A), which were higher than their MDLCs *in-vitro* counterparts (see Figure 1A). In addition, inner foreskin EpiDCs expressed CB1 and CB2 (Supplementary Figure 2), at levels that were higher compared to LCs. We also determined CB1 and CB2 expression in epidermal inner foreskin viable CD45^+^CD3^+^CD8^-^ T-cells, and found that they expressed comparable levels of both receptors (Figure 5B), which were higher compared to their blood CD4^+^ T-cell counterparts (see Figure 3A).

**Figure 5.**
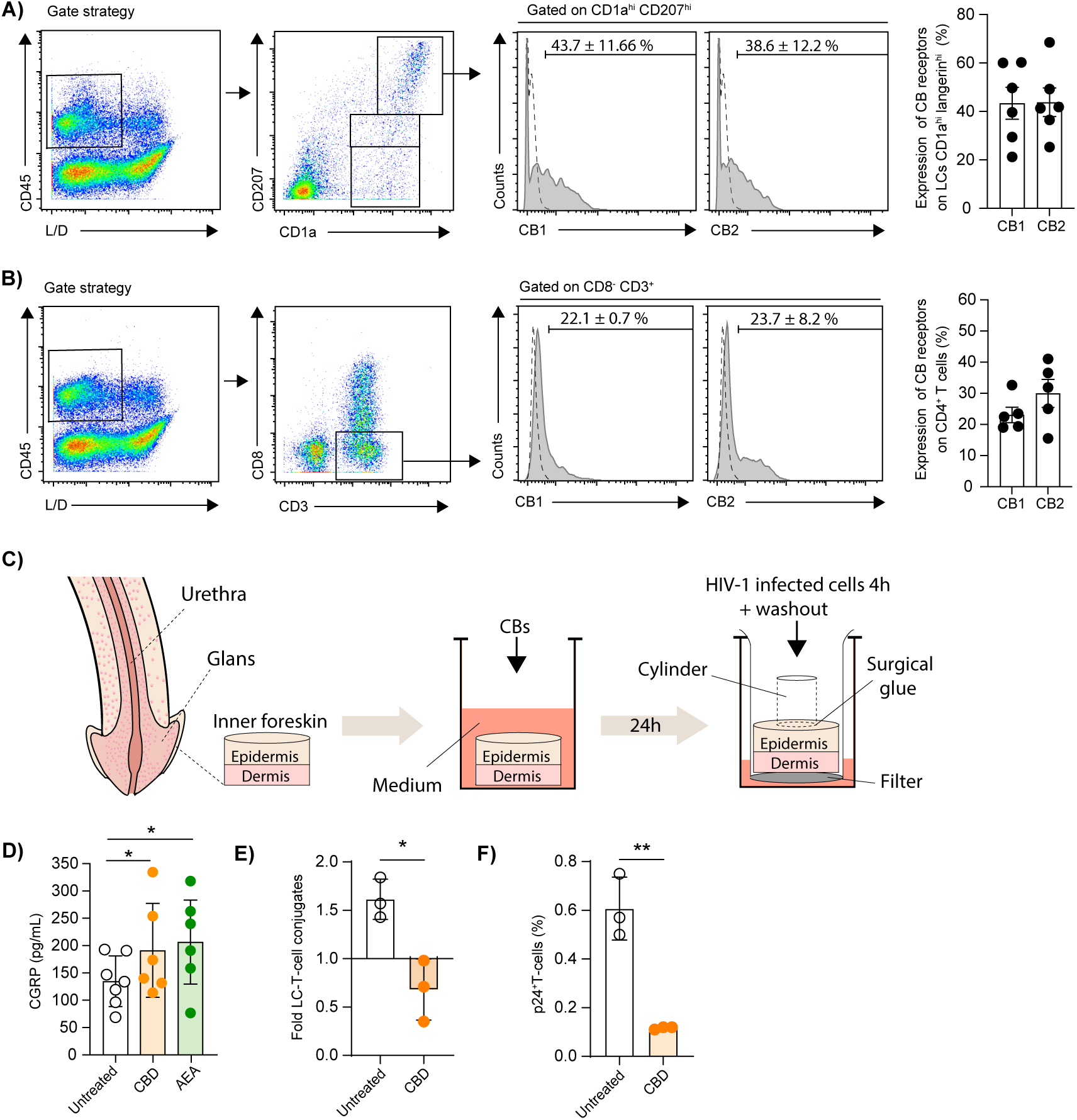
CBD induces CGRP secretion and inhibits HIV-1 R5 mediated LC-T-cells conjugates formation and CD4^+^ T-cells infection, in inner foreskin tissue explants ex-vivo. **(A, B)** Inner foreskin epidermal cell suspensions were labeled with Viobility Fixable Dye to distinguish Live/Dead (L/D) cells, stained for surface expression of CD45, CD1a/CD207 (langerin) or CD3/CD8, CB1 and CB2, and examined by flow cytometry. Shown are representative dot plots of viable CD45^+^ cells subsequently separated into CD1a^hi^CD207^hi^ LCs and CD1a^low^CD207^low/neg^ EpiDCs (A) or CD3^+^CD8^-^ T-cells (B). Also shown are representative flow cytometry overlay histograms of CB1 and CB2 surface expression in CD1a^hi^CD207^hi^ LCs (A) and CD3^+^CD8^-^ T-cells (B). Numbers and graph show mean±SD percentages of CB1^+^ or CB2^+^ cells. **(C)** Schematic representation of the different experimental steps, showing treatment with CBs followed by polarized HIV-1 infection of inner foreskin tissue explants. **(D-F)** Inner foreskin tissue explants were submerged in culture media, and either left untreated or treated with CBD or AEA (10^-5^M). In (D), media were collected 24h later and the levels of secreted CGRP were determined using an EIA. Shown are mean±SD pg/ml secreted levels of CGRP. In (E, F), untreated or CBD-treated inner foreskin tissue explants were next washed, transferred to two-chamber Transwell inserts, inoculated in a polarized manner with either non-infected or HIV-1-infected cells, and 4h later the cell-associated HIV-1 inoculum was washed out. In (E), explants were immediately digested with dispase and trypsin to obtain epidermal cell suspensions. After gating out cell debris and FSC^highest^SSC^highest^ keratinocytes, cells were further gated on viable CD45^+^CD3^+^CD8^−^ T-cells, and the percentages of FSC^hi^ conjugates with CD1a^hi^langerin^hi^ LCs were determined by flow cytometry. Shown are mean±SD folds increase in conjugate formation, calculated as [(% conjugates following inoculation with HIV-1-infected PBMCs) / (% conjugates following inoculation with non-infected PBMCs). In (F), explants were further incubated for 3 additional days submerged in fresh culture media, before digestion with dispase and collagenase/DNase to obtain dermal cell suspensions. Cells were gated on viable FSC^low^SSC^low^ lymphocytes, and the percentages of CD3^+^CD8^-^p24^+^ cells were determined by flow cytometry. Shown are mean±SD percentages of HIV-1-infected T-cells, calculated as [(%CD3^+^CD8^-^p24^+^ cells following inoculation with HIV-1-infected PBMCs) - (%CD3^+^CD8^-^p24^+^ cells following inoculation with non-infected PBMCs)]. In all graphs, dots denote different experiments using tissues from different human donors; *p<0.05, **p<0.005 and ***p<0.0005, Student’s t-test.

Next, we treated inner foreskin tissue explants with 10^-5^M CBD or AEA for 24h, and determined by an enzyme immunoassay (EIA) the levels of CGRP secreted into the media (see the scheme of experimental steps in Figure 5C). We found that both CBD and AEA treatment significantly increased CGRP secretion by these inner foreskin tissues (Figure 5D).

Finally, we used our established model of HIV-1 infection *ex-vivo* (Figure 5C), in which inner foreskin tissue explants are placed onto two-chamber inserts and inoculated in a polarized manner with cell-associated HIV-1 (i.e., PBMCs infected with an HIV-1 R5 primary isolate). We reported that such infection (but not that will cell-free virus ^1, 2^) increases formation of epidermal LC-T-cell conjugates, which occurs already and the end of the polarized infection period and can be detected by flow cytometry in epidermal cell suspensions. Of note, we recently showed that in the inner foreskin, increased conjugates formation involves LCs but not EpiDCs ^21^. In turn, these conjugates facilitate HIV-1 transfer to and infection of CD4^+^ T-cells, which can subsequently be detected by flow cytometry in dermal cell suspensions; both events are inhibited by CGRP ^15, 19, 21^ and CP ^21^.

Herein, we found that CBD pre-treatment significantly and completely abrogated the increase in LC-T-cell conjugates formation, and actually reduced the baseline levels of conjugates detected in untreated explants (Figure 5E). Importantly, CBD also blocked almost completely, namely by approximately 80%, the resulting infection of CD4^+^ T-cells (Figure 5F).

These results show that CBD inhibits the initial events of HIV-1 transmission within inner foreskin mucosal tissues *ex-vivo*.

## Discussion

CBs have been used for centuries for recreation and medicinal purposes, and medical cannabis refers to the use of marijuana-derived products prescribed by clinicians to manage symptoms of various conditions. Emerging data, along with common belief by the general population that CBD is beneficial, place CBD as a promising therapeutic compound for alleviating disease symptoms ^53^. Yet, its approval as a product for medical purposes remains restricted and, to date, only two CBD-based therapeutics are approved for clinical use, i.e., for management of refractory epilepsy (Epidiolex®) and spasticity/pain in multiple sclerosis (Sativex®) ^53^. CBD remains less rigorously studied than conventional medicines, and scientific data showing its efficacy are often missing.

As CBs are immunosuppressive, dampen pro-inflammatory pathways and shift immune tone towards resolution, previous studies investigated their role during HIV-1 infection. Some studies in PLWH concluded that cannabis use and administration are associated with reduced inflammation markers ^30, 31^, although in a recent cohort CBD was found to have no significant effects on PLWH health-related quality of life ^54^. In addition, other studies reported on inhibitory effects mainly of THC and CB1/CB2 agonists during HIV-1 infection *in-vitro* ^32–35^, while systematic evaluation of the potential anti-HIV-1 roles of CBD, across different HIV-1 immune target cells, was not performed. Herein, we reveal that CBD is endowed with important and direct anti-HIV-1 effects, by acting on all the principal mucosal HIV-1 cellular targets, namely LCs, DCs, macrophages, and importantly CD4^+^ T-cells, via different mechanisms.

In LCs and macrophages, which are ontogenetically related, CBD inhibits LCs-mediated HIV-1 transfer and macrophages HIV-1 direct infection. These effects result from preferential activation of TRPV1 and subsequent secretion of CGRP that exerts these anti-HIV-1 effects, which we previously discovered. Indeed, addition of a CGRP receptor antagonist before CBD treatment abrogates approximately half of CBD-mediated inhibition, indicating that the TRPV1/CGRP pathway plays a major role in these cells. The eCB AEA acts similarly to CBD, and we show that AEA-mediated inhibition of HIV-1 transfer is completely abrogated in the presence of a TRPV1 antagonist. Moreover, AEA-mediated inhibition HIV-1 transfer by LCs and infection of macrophages is abrogated by half in the presence of a CGRP receptor antagonist. While the remaining inhibitory effects of CBD might be attributed to CB2 activation in macrophages, this is not the case in LCs in which the CB1/CB2 agonists show no effects. Further studies in LCs should now determine which other receptors that CBD was reported to activate (e.g., peroxisome proliferator-activated receptor gamma (PPARγ) or the serotonin 1A receptor (5-HT_1A_)) ^26^ could be involved.

We further show that in LCs, CBD and AEA inhibit the late phase of HIV-1 transfer to CD4^+^ T-cells, which involves the transfer of *de-novo* produced virions. Accordingly, CBD and AEA could mediate this effect via the downregulation of langerin/CD4/CCR5 expression in LCs that we observed, which would limit LCs productive HIV-1 R5 infection and hence late transfer. Noteworthy is the production of eCBs by LCs and their downregulation upon HIV-1 infection, indicating the existence of a potential endogenous eCB-mediated anti-HIV-1 mechanism in LCs that is counteracted by the virus. It would be of interest now to determine whether other eCBs, besides AEA, modulate HIV-1 infection.

In contrast, in DCs and CD4^+^ T-cells, inhibition of HIV-1 transfer and/or infection is CGRP-independent, in line with our previous observations that CGRP does not affect DCs-mediated HIV-1 transfer ^15^ or direct infection of CD4^+^ T-cells ^21^. We speculate nevertheless that TRPV1 activation in T-cells is the underlying mechanism of CBD-mediated inhibition, as we recently reported that CP, the natural TRPV1 agonist, also inhibits T-cell infection with HIV-1 ^21^. CBD-mediated inhibitory effects in T-cells could be related to downregulation of T-cell activation and CD4/CCR5 expression, as we show herein. Interestingly, the functional effects of AEA and CBD differ, as AEA fails to modulate T-cell activation and CD4/CCR5 expression. Hence, the inhibitory effects of AEA are restricted to HIV-1 X4 viruses, and could be mediated by its reported capacity to activate CB2 (that we confirmed) and inhibit CXCR4-medited signaling ^48^. Importantly, we show that our *in-vitro* findings translate to efficient control of mucosal HIV-1 transmission *ex-vivo*. In our model of polarized infection of inner foreskin tissue explants with HIV-1-infected cells, we found that pre-treatment with CDB induces CGRP secretion from the explants, and is accompanied by inhibition of two major transmission parameters. These include: i) complete block, and even reversal, of HIV-1-induced increased formation of epidermal LC-T-cell conjugates, immediately following HIV-1 inoculation; ii) significant inhibition by approximately 80% of the later ensuing HIV-1 infection of CD4^+^ T-cells. These findings provide ‘proof-of-concept’ for the efficacy and of CBD in preventing mucosal HIV-1 transmission. Despite the existence of both prophylactic (pre-exposure prophylaxis - PrEP, and post-exposure prophylaxis - PEP) and therapeutic ART strategies, HIV-1 remains a global public health concern. PrEP consisting of the antiretrovirals tenofovir disoproxil / emtricitabine is available and highly effective for prevention of HIV-1 transmission, and recent major advancements showed the unprecedent efficacy of long-lasting injectable cabotegravir and lenacapavir. The question hence arises: is HIV-1 prevention research still needed? We argue that this is undoubtedly the case, as two central issues still limit PrEP implementation: i) choice - as recently concluded, offering a variety of adaptable prevention methods, in addition to ‘classical’ PrEP, ensures optimal adherence to prevention efforts, as individuals can make informed decisions that best suit their circumstances ^55^; ii) limiting barriers - cost, access, adherence, stigma, adverse side effects and eventual drug resistance ^56^. Moreover, 3.5 million persons globally initiated or continued PrEP during 2023, far fewer than the United Nations Program on HIV/AIDS goal of at least 21.2 million persons initiating or continuing PrEP globally in 2025. This gap highlights the ongoing need not only for investment in expanding PrEP access worldwide ^57^, but also for innovation and development of other prevention strategies.

Bases on our results, we propose an alternative that we coin as ‘CBD PrEP’. This would consist of the repositioning of commercially available CBD-containing formulations as novel microbicides for the purpose of clinical HIV-1 prevention. Future studies should now evaluate the efficacy, for instance of already formulated lubricants that contain CBD and are intended for topical use, for this purpose (i.e., in which even as low as 1% CBD equals 30mM that is above the highest concentrations tested herein, which could ensure penetration of effective CBD concentrations into mucosal tissues upon topical application). These CBD-based products would have several advantages over existing PrEP: i) cost - cheap; ii) accessibility - available to the general public in ‘CBD shops’ and pharmacies; iii) stigma - popular, on high-demand and well accepted for their benefits by the general population; iv) adverse effects - safe and devoid of side effects / drug resistance. Our CBD-based approach is an original strategy for HIV-1 prevention, which could complement current measures that remain inadequate, insufficient and not optimal. Even if exerting only partial protection against HIV-1 acquisition, CBD PrEP could contribute to a global reduction in HIV-1 burden, especially in middle- and low-income countries where preventive measures are the most needed.

## Acknowledgments

The study was funded by SIDACTION, via a research grant to YG (N/Ref 19-2-AEQ-12566) and a post-doctoral fellowship to CCBB (AAP31-2-FJC-12895). DC and IM acknowledge INSERM, Nouvelle Aquitaine Region and University of Bordeaux IdEx “Investments for the Future” program/GPR BRAIN_2030.

## Author Contributions

CCBB and YG designed research; CCBB, HG, JM, IM, DC and YG performed research and analyzed data; NBD and MZ contributed new reagents/analytic tools; CCBB, MB and YG wrote and edited the paper.

