## Supplementary material for "Cannabidiol prevents mucosal HIV-1 transmission by targeting Langerhans cells, dendritic cells, macrophages and T-cells": Supp figures

#### Supplementary Figure legends

**Supplementary Figure 1. CB2 intracellular expression in human LCs, $CD4^+$ T-cells and macrophages.** Representative flow cytometry overlay histograms showing CB2 intracellular expression in MDLCs, gated on $langerin^+$ cells **(A)**, primary blood $CD4^+$ T-cells **(B)**, and MDMs **(C)**. Numbers and graphs show mean±SD percentages of $CB2^+$ cells.

**Supplementary Figure 2. CB1 and CB2 surface expression in human inner foreskin EpiDCs.** Representative flow cytometry overlay histograms showing CB1 and CB2 surface expression in $CD1a^{low}CD207^{low}$ (upper panel) and $CD1a^{low}CD207^{neg}$ (bottom panel) inner foreskin EpiDCs. Numbers and graphs show mean±SD percentages of $CB1^+$ and $CB2^+$ cells.

### Supplementary figures

#### Supplementary figure 1

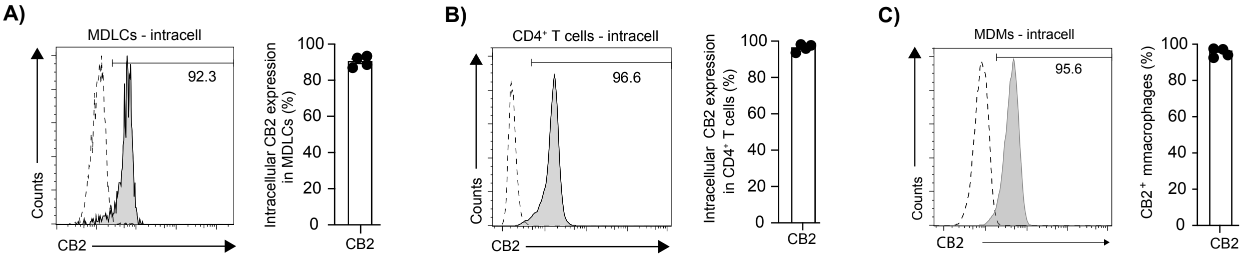


#### Supplementary figure 2

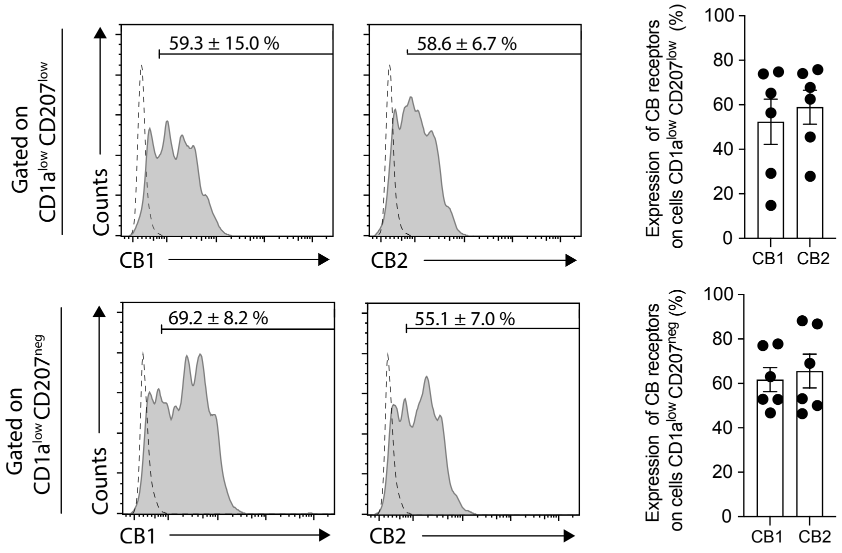
